# cNap1 bridges centriole contact sites to maintain centrosome cohesion

**DOI:** 10.1101/2021.11.20.469313

**Authors:** Robert Mahen

## Abstract

Centrioles are non-membrane bound organelles that participate in fundamental cellular processes through their ability to form physical contacts with other structures. During interphase, two mature centrioles can associate to form a single centrosome - a phenomenon known as centrosome cohesion. Centrosome cohesion is important for processes such as cell migration, and yet how it is maintained is unclear. Current models indicate that pericentriolar fibres termed rootlets, also known as the centrosome linker, entangle to maintain centriole proximity. Here, I uncover a new centriole-centriole contact site and mechanism of centrosome cohesion, based on coalescence of the proximal centriole component cNap1. Using live-cell imaging of endogenously tagged cNap1, I show that proximal centrioles form dynamic contacts in response to physical force from the cytoskeleton. Expansion microscopy reveals that cNap1 bridges between these contact sites, physically linking proximal centrioles on the nanoscale. When ectopically tethered to organelles such as lysosomes, cNap1 forms viscous and cohesive condensates that promote organelle spatial proximity. Conversely, cNap1 mutants with reduced viscosity are unable to maintain centrosome cohesion. These results define a previously unrecognised mechanism of centrosome cohesion by cNap1 assemblies at the proximal centriole and illustrate how a non-membrane bound organelle forms dynamic organelle contact sites.

## Introduction

Organelle contact sites are critical to diverse cellular functions. Membrane bound organelles associate via dedicated molecular complexes that perform functions such as membrane tethering [1]. How non-membrane bound molecular assemblies form physical contacts with other cellular structures is less clear.

Centrosomes are microtubule organising centres that mediate fundamental cellular processes including cell division, polarity and motility. Centrosomes exist in the cellular interior without a bounding membrane, dynamically interacting with structures such as the cell membrane and mitotic spindle [2]. During interphase, mammalian centrosomes contain two mature microtubule-based structures called centrioles. Centrioles associate together in a process termed centrosome cohesion [3,4]. Centrosome cohesion is important for mitosis, ciliary function and cell migration [5–8], and thus understanding its molecular and biophysical basis is an important question.

Rootlets, also known as the centrosome linker, are pericentriolar fibres found at centrioles [3,9,10]. Across the Animalia kingdom, rootletin protein (*CROCC*) is a key component of rootlets [5,11–14]. Rootlets are frequently prominent when centrioles form cilia in specialised cell types such as mechanosensory neurons, and may form links between centrioles as part of polarised multiciliary arrays [10]. Rootletin loss of function studies have demonstrated that rootlets are required for centrosome cohesion in non-ciliated human cells [3,15,16]. One model postulates that rootlets maintain centrosome cohesion by entangling together, therefore establishing direct physical links between centrioles [3,9,17–19]. These links are likely weak or dynamic, since centrioles at times transiently separate, potentially in response to physical force from the cytoskeleton [20–24].

cNap1 (also known as CEP250) is a rootletin paralog found at the proximal centriole, spatially adjacent to rootlet fibres [18,19,25,26]. Truncating mutations in cNap1 cause mammalian retinal and developmental dysfunction [7,27,28]. cNap1 binds to rootletin in biochemical assays, dissociates from centrosomes when they split during mitosis, and is required for rootlet formation at centrosomes, suggesting that it anchors rootlets to centrioles [3,9,15,26,29]. cNap1 disruption also causes loss of centrosome cohesion [7,15,29,30] – a phenomenon attributed to rootlet disruption. Little is known about the biophysical properties of cNap1 that allow it to maintain centrosome cohesion, or the molecular basis of centriole-centriole contact sites.

Here, by studying the biophysical properties and nanoscale architecture of cNap1 at centriole-centriole contact sites, I discover that it directly maintains centrosome cohesion. Live-cell imaging and expansion microscopy of endogenous cNap1 shows that it bridges between dynamic centriole-centriole interfaces. Overexpression of cNap1 forms biomolecular condensates with viscous material properties that cohesively maintain organelle spatial proximity. Conversely, mutants with reduced cNap1 viscosity fail to maintain centrosome cohesion. I propose a model of centrosome cohesion explaining how organelle solidity is balanced against organelle plasticity using dynamic cNap1 assemblies.

## Results

### Proximal centriole pairs and rootlets form dynamic contacts during centrosome cohesion

To simultaneously track the spatiotemporal behaviour of both proximal centrioles and rootlets in living cells, genome editing was used to create cells expressing endogenously tagged fluorescent cNap1 and rootletin (cNap1-mScarlet-I and rootletin-meGFP). Cell lines were carefully validated; precise genomic tagging was ensured by a combination of overlapping genomic PCR and imaging (**S1A-C Fig**. and **Materials** and **methods**). Clones were screened to identify cells with all cNap1 alleles homozygously modified. cNap1-mScarlet-I and rootletin-meGFP functionality was ensured by measuring centrosome cohesion, which was indistinguishable in genome edited and wild type cells (**S1D Fig**.). These considerations confirmed that cNap1 and rootletin were tagged at functional, endogenous levels.

I reasoned that if centrosome cohesion is mediated by direct links between centrioles, these contact points might be visible with imaging. cNap1-mScarlet-I enriched strongly at centrosomes relative the surrounding cytosolic pool, forming either one or two foci corresponding to the proximal centrioles (**Fig. 1A**), as expected from prior electron microscopy [26]. Live-cell Airyscan time-lapse imaging showed that on the seconds to mins timescale, the relative spatial proximity of centriole pairs was variable (**Fig. 1B** and **Movie 1**). Proximal centrioles thus transiently formed contacts as they collided together during continuous movement. Rootlet fibres were also mobile during centriolar movements. Rootlets from both centrioles could either form contacts, or alternatively move independently, apparently not in contact (**Fig. 1C** and **1D; Movies 2** and **3**). Throughout these dynamic movements, cNap1-mScarlet-I foci were always present at the centrosome-proximal termini of rootletin-meGFP fibres (**Fig. 1E**). Thus, simultaneous imaging of endogenously tagged cNap1 and rootletin reveals that both proximal centrioles and rootlets form dynamic contacts during centrosome cohesion.

**Fig 1.**
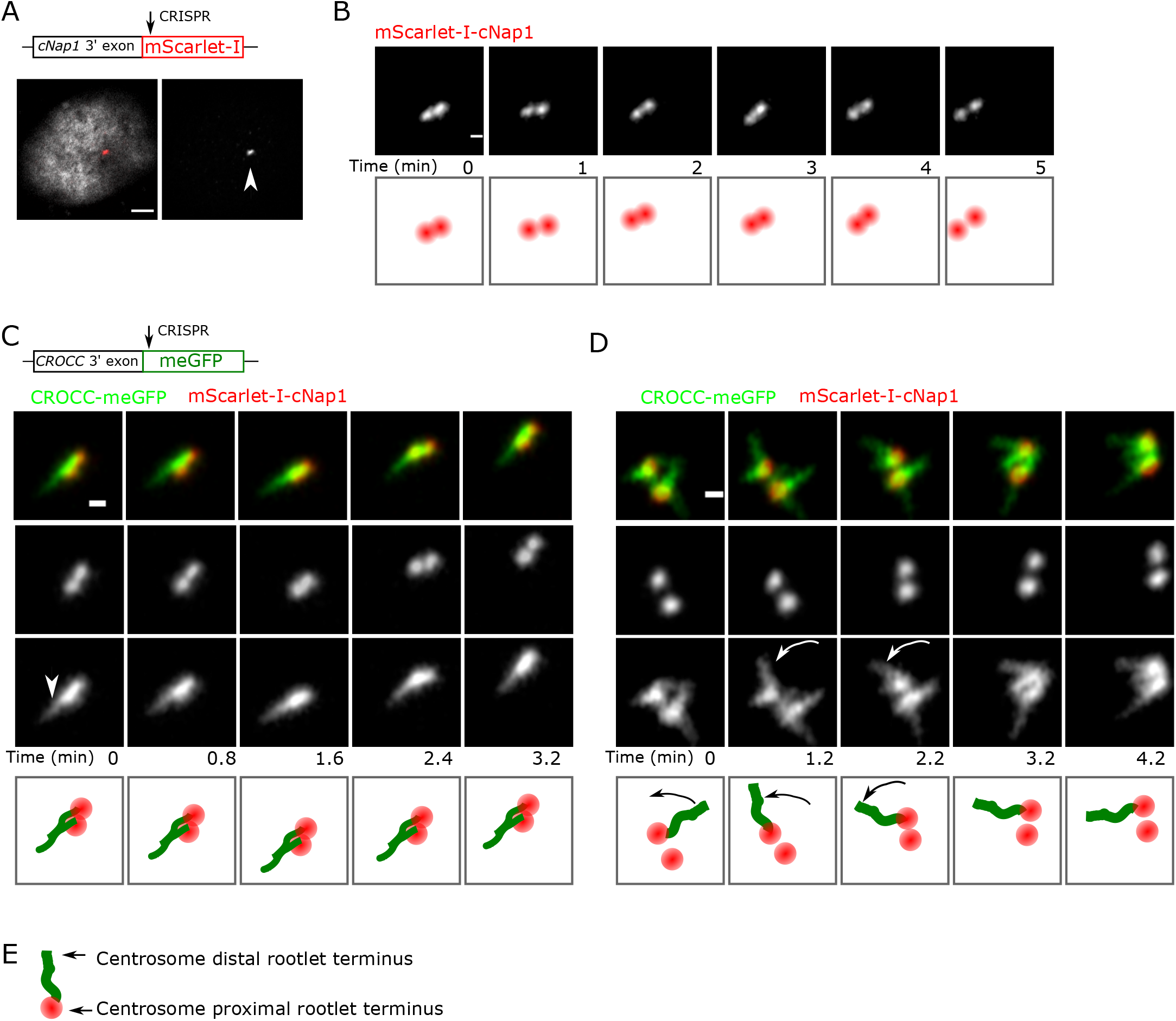
Proximal centriole pairs and rootlets form dynamic contacts during centrosome cohesion. (**A**) Endogenously tagged cNap1-mScarlet-I localises to regions of high concentration at proximal centrioles (denoted by the arrow). A merged image showing both cNap1-mScarlet-I and the nucleus is shown on the left panel. The right panel shows only cNap1-mScarlet-I. Scale 3 μm. (**B**) Time-lapse imaging of endogenously tagged cNap1-mScarlet-I at one-min intervals shows dynamic contacts. The images show maximum intensity projections from 3D data. Scale 0.5 μm. (**C-D**) Time-lapse two-colour Airyscan imaging of endogenously tagged cNap1-mScarlet-I and rootletin-meGFP. Scale 0.5 μm. The arrowhead in (**C**) denotes a potential point of contact between rootlets from different centrioles. The arrows in (**D**) denote independent movement of a rootlet distal terminus relative to other rootlets. (E) Cartoon depiction of the arrangement of the paralogs cNap1 and rootletin at centrosomes; centrosome proximal cNap1-mScarlet-I is attached to rootlet termini.

### Endogenous cNap1 bridges proximal centrioles at the nanoscale

To investigate centriole-centriole contact sites below the diffraction limit of light microscopy, I turned to ultra-expansion microscopy (U-ExM) [31]. U-ExM increases resolution by expanding fixed cells approximately fourfold in size, therefore achieving resolution on the tens of nanometres scale [31]. Since centriole diameter and length have previously been determined, I used anti-acetylated tubulin staining of centrioles to ensure isotropic and efficient expansion in my experimental setup, with previously described protocols optimised to maintain centriole morphology [31,32]. Anti-acetylated tubulin staining of centriolar barrels gave a diameter of ~190 nm, consistent with previous work, and demonstrating isotropic and efficient expansion [32].

Consistent with the live-cell imaging data (**Fig. 1**), centriole pairs occupied variable orientations relative to each other in populations of cells (**Fig. 2A)**. U-ExM staining of cNap1 with an siRNA-validated antibody (**S2A Fig**.), showed that it accumulates at the proximal centriole as expected [26]. Strikingly, cNap1 formed structures that bridged both centrioles in 39% of cells (from a total of 80 cells imaged). Thus, cNap1 from both centrioles either merged together into a single ellipsoid-shaped structure, or two ellipsoids could form junctions between two centrioles in various orientations. In the remaining 61% of cells, cNap1 formed two unconnected spatially separate structures. Imaging end-on down the centriole barrel highlighted variability in the cNap1 structures as asymmetrical ellipsoids (**Fig. 2B**). Regardless of whether centrioles were cohered or not, cNap1 was frequently located spanning from inside the centriole barrel to outside it (**Fig. 2C)**. A montage of ~60 different cells is presented in **S2B Fig**. to document this variability in cNap1 orientation and shape.

**Fig 2.**
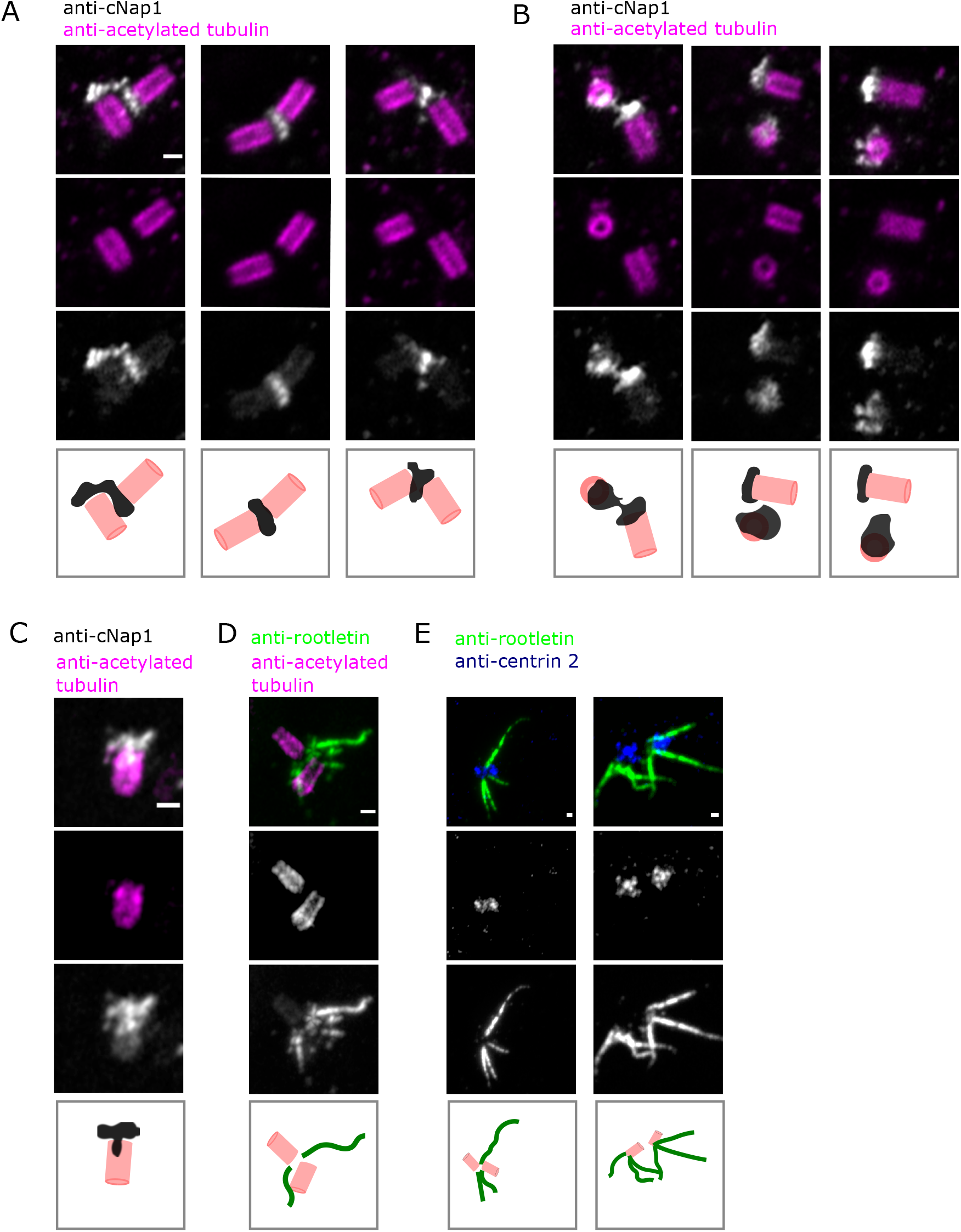
Endogenous cNap1 bridges proximal centrioles at the nanoscale. (**A-C**) U-ExM of centrioles labelled with anti-acetylated tubulin antibody (pink), and cNap1 labelled with anti-cNap1 antibody (grey). The images show z-slices. Cartoons depict simplified centriole and cNap1 orientations. (**D**) U-ExM of rootlets stained with anti-rootletin antibody (green) and centrioles stained with anti-acetylated tubulin antibody (pink). (**E**) U-ExM of rootlets stained with anti-rootletin antibody (green) and centrioles stained with centrin-2 antibody (blue). Maximum intensity projections are shown. Across all panels the scale is 200 nm, and each column represents a different cell.

U-ExM of rootletin showed that it too was present in multiple orientations, also consistent with the live-cell imaging data (**Fig. 2D**). With centrioles adjacent, rootlets could either be oriented radially into the cytoplasm without overlap, or formed contact points with the rootlets from the adjacent centriole (**Fig. 2E**). Together these results demonstrate that cNap1 can either form separate assemblies on each proximal centriole or can bridge centriole pairs as a contiguous structure, therefore merging both proximal centrioles.

### cNap1 forms viscous condensates

I investigated further the biophysical properties of cNap1 that allow it to bridge across centriole pairs. Overexpression of mScarlet-I-cNap1 resulted in cytosolic patches in various mammalian cell types, in agreement with previous studies [9,33,34]. Thus, at low expression levels mScarlet-I-cNap1 localised to the centrosome as seen by co-staining with the centrosomal marker gamma-tubulin, and at higher expression levels mScarlet-I-cNap1 additionally formed cytoplasmic patches of variable size and number (**Fig. 3A**). mScarlet-I-cNap1 patches were ellipsoid in shape, becoming more elongated at larger sizes as measured by automated imaging and analysis in populations of cells, and formed spontaneously over a threshold concentration (**Movie 4**). Many cellular structures have recently been posited to be biomolecular condensates, with material properties equivalent to states of matter such as solids or liquids [35,36]. I therefore investigated the material properties of mScarlet-I-cNap1 in cytosolic patches using fluorescence recovery after photobleaching (FRAP) and live single-cell imaging. Over short timescales of up to ~30 s, mScarlet-I-cNap1 showed little diffusional mobility, indicating limited internal rearrangement of component parts on this timescale (**Fig. 3B)**, consistent with previous data [33]). However, over longer timescales of mins to hours, mScarlet-I-cNap1 patches showed viscous liquid-like behaviour and were able to coalesce (**Fig. 3C**; **Movie 5** and **6**: **S3A Fig**.). During coalescence, mScarlet-I-cNap1 patches first retained their shape whilst touching, before slowly forming intermediate bridged shapes and then one contiguous structure. These observations are consistent with mScarlet-I-cNap1 forming viscous condensates with a propensity to coalesce.

**Fig 3.**
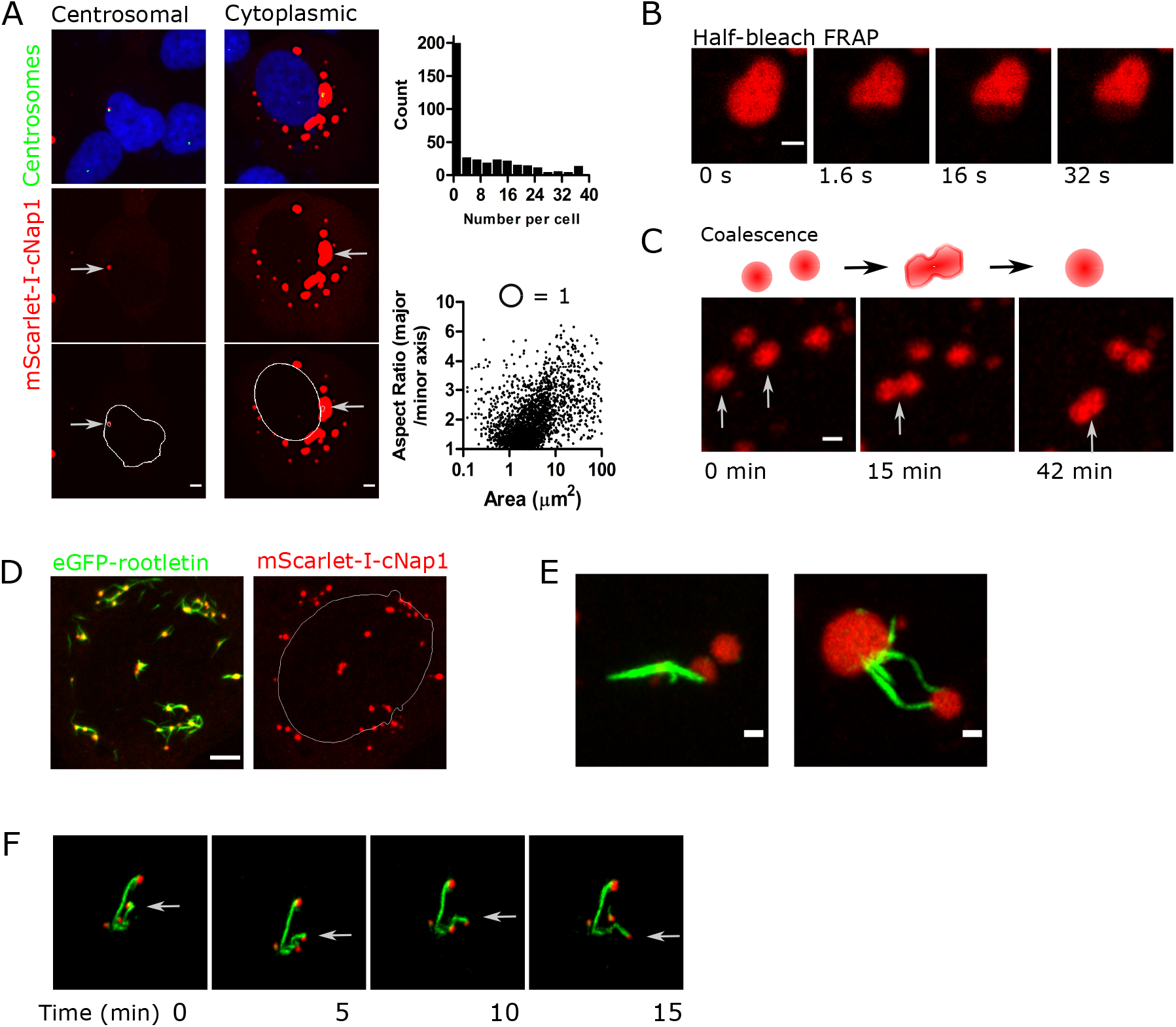
cNap1 forms viscous condensates that bind rootletin fibre termini. (**A**) Ectopic cNap1-mScarlet-I expression results in either centrosomal or cytosolic patches (red). Centrosomes are co-stained with gamma-tubulin (green), and centrosome position is indicated with arrows. White lines in the bottom panels denote nuclei. Scale bar 4 μm. The histogram shows the number of mScarlet-I-cNap1 patches per cell in a population of 388 cells, acquired with automated imaging and analysis as detailed in Materials and methods. The x, y graph plots mScarlet-I-cNap1 area against aspect ratio (long axis / short axis) in ~4000 patches, where a circle has an aspect ratio of 1. (**B**) Half-bleaching FRAP of an mScarlet-I-cNap1 patch in the cytosol shows limited exchange over ~30 s. The bleached region is located at the bottom, and images show successive indicated timepoints. Scale bar 2 μm. (**C**) Live-cell time-lapse imaging of mScarlet-I-cNap1, showing coalescence of a viscous liquid. Maximum intensity projections are shown. Scale 1 μm. (**D**) Co-expression of mScarlet-I-cNap1 (red) and eGFP-rootletin (green) in a single cell. Scale 5 μm. The white line indicates the location of the nucleus for reference. (**E)** Two detailed views of mScarlet-I-cNap1 condensates associated with eGFP-rootletin fibres. Left panel: mScarlet-I-cNap1 at a single eGFP-rootletin fibre terminus. Right panel: mScarlet-I-cNap1 at both eGFP-rootletin termini. Scale bars 1 μm. (**F**) Co-movement of mScarlet-I-cNap1 condensates and rootletin-meGFP fibre termini (denoted by the arrow) in live-cell time-lapse imaging over a period of mins.

### cNap1 condensates bind rootletin fibre termini

cNap1 has previously been shown to bind rootletin by yeast two-hybrid and coimmunoprecipitation [3,9]. I therefore investigated whether rootletin is recruited into mScarlet-I-cNap1 condensates. mScarlet-I-cNap1 was co-expressed in cells stably expressing eGFP-rootletin, to allow dual-colour time-lapse imaging of the spatiotemporal behaviour of both transgenes during condensate formation. Surprisingly, eGFP-rootletin was not recruited into cNap1 condensates per se. Instead, mScarlet-I-cNap1 condensates bound to eGFP-rootletin fibre termini (**Fig. 3D** and **Movie 7**). Thus, ~63% of eGFP-rootletin fibres were coincident with mScarlet-I-cNap1 patches, ~18 hours after mScarlet-I-cNap1 transfection in a population of cells. cNap1 predominantly bound to the ends of cytoplasmic rootletin fibres, attaching to either one or both ends (**Fig. 3E**). mScarlet-I-cNap1 was not just temporarily colocalised with rootletin fibre termini, but stably attached, since they could translocate together over mins (**Fig. 3F** and **Movie 8**). Thus, mScarlet-I-cNap1 binds to rootletin fibre termini when overexpressed, self-organising into the same spatial arrangement as endogenous cNap1 foci at centrosomes (**Fig. 1**). Moreover, these results demonstrate that cNap1 forms condensates with viscous material properties.

### cNap1 condensate formation promotes rootlet end-binding but not centrosomal localisation

To investigate whether individual cNap1 protein domains are sufficient for condensate formation and protein viscousity, cNap1 was divided into a series of separate fragments as described previously [37]. These protein fragments consist of either the N terminus, the C terminus, the middle domain 1, the middle domain 2 or both middle domains (**Fig. 4A** and **Supplementary Table 1;** respectively named here; NT, CT, M1, M2 or M1/2). An R188 mScarlet-I-cNap1 truncation was also created, since it is the site of a truncating mutation (R188) identified in a consanguineous family with retinal impairment [27,28], and close to a nearby truncating reside (169) reported to cause developmental defects in cows [7].

**Fig 4.**
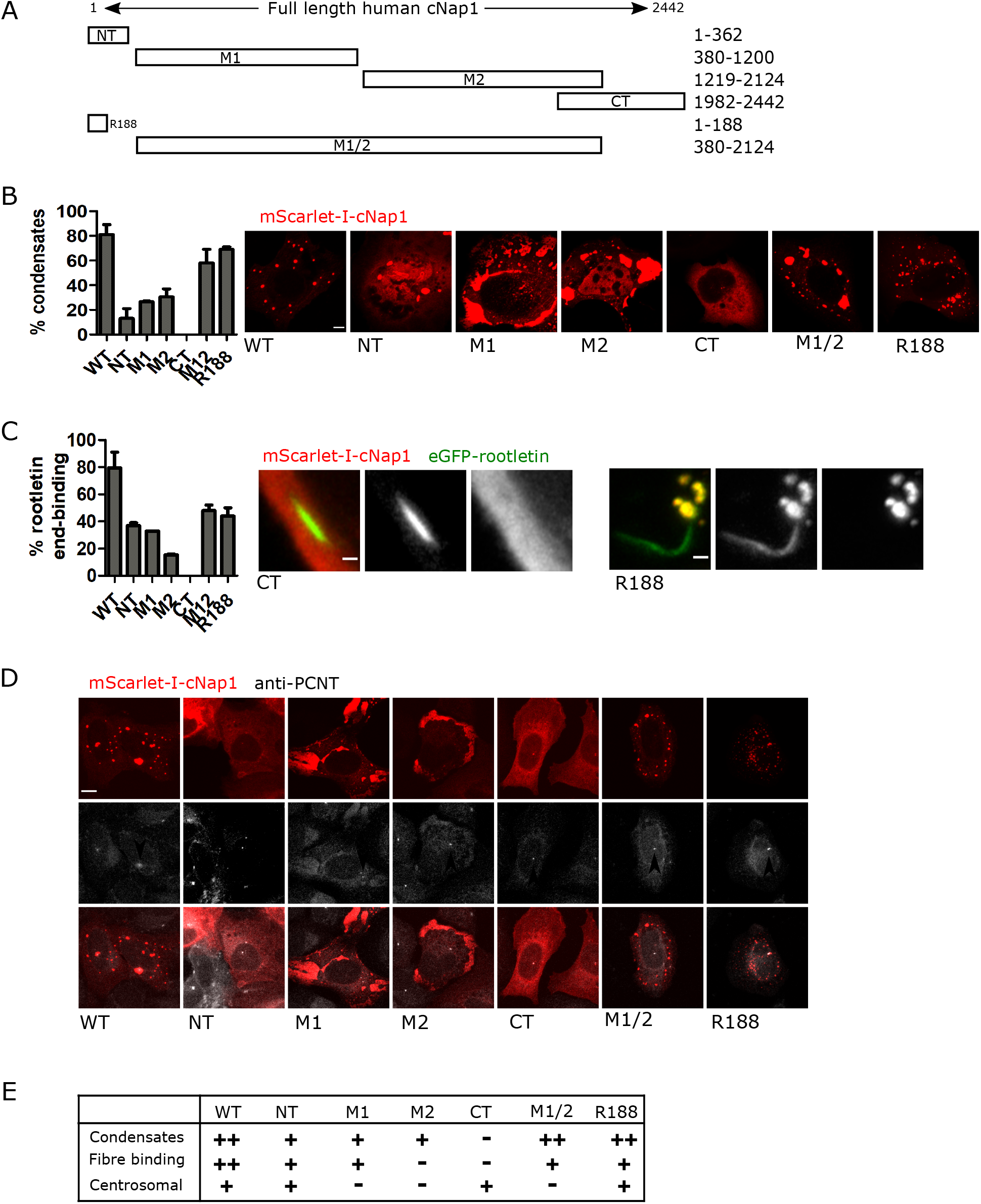
cNap1 condensate formation promotes rootlet end-binding but not centrosomal localisation. (**A**) Schematic representation of cNap1 protein truncations. Numbers denote amino acids from the N-terminus. (**B**) Condensate formation by cNap1 truncations. The bar graph plots the mean percentage of cells containing greater than one condensate, from two experiments. The images show a representative example cell with condensates (except CT). Scale 5 μm. (**C**) Rootletin fibre binding by cNap1 truncations. The bar graph plots the percentage of rootletin fibres associated with cNap1 condensates, from two experiments. Cells without condensates were excluded from the analysis. The images show a representative rootletin fibre (green) with cNap1 CT or R188 (red). Scale 10 μm. (**D**) Centrosomal localisation by cNap1 truncations. The representative confocal images show mScarlet-I-cNap1 (red) and centrosomes marked by anti-PCNT (white). Centrosomes are indicated by the arrows. Maximum intensity projections are shown. The “smooth” function was used in Fiji and image brightness and contrast are changed for display purposes. Scale 10 μm. (**E**) Summary of mScarlet-I-cNap1 truncation properties, from the experiments in (B - D). ++, + and – denote decreasing amounts respectively.

No single cNap1 protein domain was fully sufficient for condensate formation at wild type levels (**Fig. 4B**). Instead, dependent on the domain, 0-70 % of cells formed cytoplasmic condensates, relative to 80 % in wild type. I investigated whether condensate formation influenced cNap1 rootlet-end binding. Rootletin fibre end-binding correlated with condensate formation capability, when analysing only cells that formed condensates. Wild type thus had the highest rootletin fibre end-binding, and other truncations had lower levels (**Fig. 4C)**. Importantly, and in contrast to rootletin fibre end-binding, condensate formation was not essential for centrosomal localisation (**Fig. 4D**). Thus, in agreement with previous work [15], either of the terminal domains (NT or CT) were sufficient for centrosomal localisation, as was R188, despite differing condensate forming ability (**Fig. 4E**). These results together suggest that condensate formation promotes cNap1-rootletin fibre end-binding but is not essential for centriolar localisation.

### cNap1 promotes organelle cohesion through its viscosity

Previous loss of function studies have shown that cNap1 is required for centrosome cohesion [7,15,29,30]. mScarlet-I-cNap1 forms condensates that coalesce (**Figs. 1** and **3**), suggesting that cNap1 itself possesses cohesive properties that might promote organelle cohesion. Such a model theorises that separate cNap1 pools coalesce to maintain spatial proximity (**Fig. 5A**). Since rootlets are already known to promote centrosome cohesion [3], this hypothesis was tested by targeting cNap1 ectopically to cellular structures not containing rootlets. Three different mScarlet-I-cNap1 fusion proteins were created, targeting mScarlet-I-cNap1 condensates to lysosomes, the Golgi and mitochondria. These constructs are termed lyso-cNap1, Golgi-cNap1 and mito-cNap1 (see **Materials and methods** for details). Note that cNap1 has not been reported to localise to these organelles. Lyso-cNap1, Golgi-cNap1 and mito-cNap1 had different shapes, related to the structure and dynamics of the targeted organelles. Lyso-cNap1 formed spherical shells surrounding lysosomes marked by LysoTracker (**Fig. 5B**). MitocNap1 and Golgi-cNap1 formed elongated structures adjacent to or coincident with either mitochondria or the Golgi respectively (**Fig. 5C** and **5D**), demonstrating that the shape of cNap1 condensates changes when targeted to different organelles.

**Fig 5.**
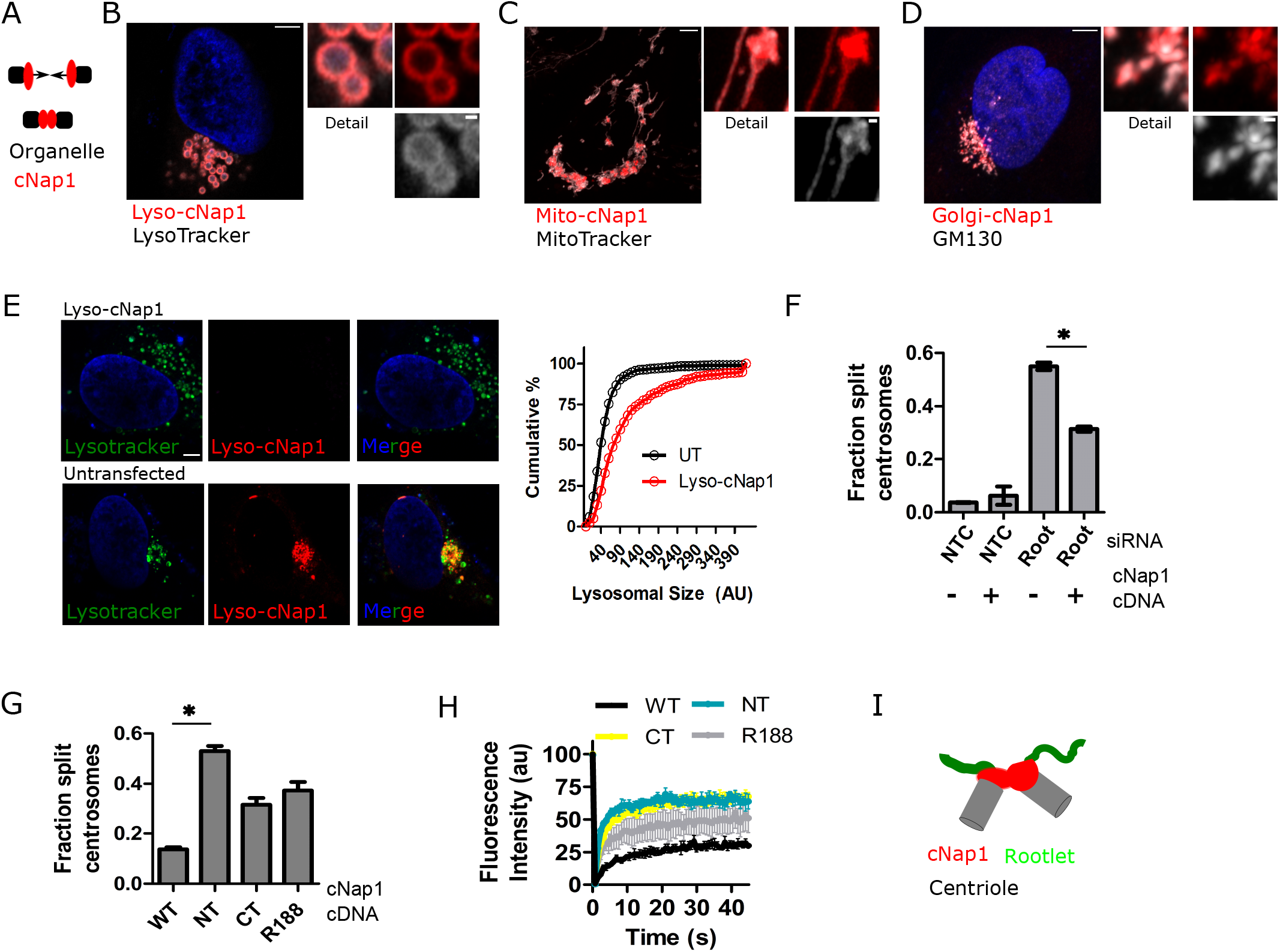
cNap1 promotes organelle cohesion through its viscosity. (**A**) Theory that organelle bound cNap1 promotes organelle spatial proximity. (**B**) Lyso-cNap1 (red) forms spherical structures coating LysoTracker positive vesicles (grey). The image shows an Airyscan confocal z-slice. Scale in large image 5 μm. Detail scale 0.5 μm. (**C**) Mito-cNap1 (red) localises adjacent to mitochondria as marked by MitoTracker (grey). Scale in large image 5 μm. Detail scale 0.5 μm. (**D**) Golgi-cNap1 (red) is elongated in shape and localises to the Golgi, as shown by co-staining with GM130 (grey). Scale in large image 5 μm. Detail scale 0.5 μm. (**E**) LysoTracker-positive vesicle localisation in the presence or absence of lyso-cNap1. The graph quantitates vesicle size either with or without lyso-cNap1 expression. (**F**) Loss of centrosome cohesion caused by rootletin siRNA is partially rescued by mScarlet-I-cNap1. The bar graph plots the percentage of cells with centrioles separated > 1.6 μm, determined from anti-PCNT staining and confocal imaging, in two experiments. The mean and the standard deviation are shown. The asterisk denotes a significant difference by t-test. (**G**) Expression of cNap1 truncations NT, CT and R188 disrupts centrosome cohesion. The bar graph plots the percentage of cells with centrioles separated > 1.6 μm, determined from anti-PCNT staining and confocal imaging, in two experiments. The mean and the standard deviation are shown. The asterisk denotes a significant difference by paired t-test (p=0.0014). (**H**) cNap1 truncations NT, CT and R188 have an increased rate of FRAP recovery at centrosomes relative to wild type. The graph plots the mean and standard error of three separate experiments, with each experiment measuring at least ten cells. (**I**) Cartoon model of cNap1-based centrosome cohesion through contact of proximal centrioles.

Targeting cNap1 to either lysosomes, the Golgi or mitochondria promoted organelle cohesion in all cases. Whereas in control or untransfected cells, lysotracker positive vesicles were spaced out within the cytoplasm (**Fig. 5E**), lyso-cNap1 coated lysosomes grouped together, frequently forming groups cohered in a honey-comb shape, an arrangement not seen in wild type cells (**Fig. 5E**, bottom panel). Time-lapse imaging revealed that lyso-cNap1 coated lysosomes also had reduced movements relative to controls (**S3B Fig**. and **C; Movie 9** and **10**). Similarly, mito-cNap1 coated mitochondria cohered together in groups with reduced movement (**Movie 11** and **S3D Fig**.).

Given that these organelles do not contain rootlets, this suggests that cNap1 itself can mediate organelle cohesion. To test whether cNap1 condensates are also sufficient to promote centrosome cohesion, I expressed mScarlet-I-cNap1 in U2OS cells, which maintain high levels of centrosome cohesion [23,29], that can be reduced by siRNA mediated knockdown of rootletin with a previously described siRNA [3]. cNap1-mScarlet-I expression significantly increased centrosome cohesion in rootletin siRNA treated cells (**Fig. 5F**). Thus, cNap1 restrains organelle spatiotemporal movement and increases organelle-organelle interaction time when ectopically targeted.

Expression of terminal cNap1 truncations with centrosomal localisation, namely CT, NT and R188, reduced centrosome cohesion, consistent with a previous study [15]. These three cNap1 domains caused increased numbers of interphase split centrosomes in a dominant negative fashion, relative to matched wild type control (**Fig. 5G**). I reasoned that if the viscosity of cNap1 contributes to centrosome cohesion, then mutants with loss of centrosome cohesion might have altered material properties. Consistent with this hypothesis, FRAP of NT, CT, and R188, revealed increased exchange rates at centrosomes relative to wild type protein (**Fig. 5H**). Moreover, timelapse imaging of R188 condensates showed reduced fusogenic capability (**Movie 12**). Together, these results show that cNap1 condensates promote centrosome cohesion, likely through their viscous cohesive properties.

## Discussion

Current models of centrosome cohesion indicate that rootlet fibres entangle to link centrioles [3,9,18,23], balancing cytoskeleton-generated forces [20,22,38]. Collectively, the results in this study provoke the hypothesis that cNap1 also directly forms inter-centriolar linkages that maintain centrosome cohesion (**Fig 5I**). Firstly, nanoscale imaging indicates that cNap1 directly bridges between centrioles (**Figs 1** and **2**). Secondly, cNap1 forms viscous condensates that promote organelle cohesion, even in the absence of rootlets (**Figs 3** and **4**). Thirdly, cNap1 mutants with reduced viscosity disrupt centrosome cohesion (**Fig 5**).

A key feature of this model is the capability of cNap1 to form viscous oligomeric assemblies, of variable shape and with cohesive properties. There is significant flexibility in the maintenance of centrosome cohesion, with centrioles able to transiently separate [23,24]. I propose that viscous cNap1 material properties allow organelle plasticity to be balanced against solidity in response to physical force from the cytoskeleton. This model may also explain previous observations that proteins with no known role in rootlet formation are required for centrosome cohesion [17].

In contrast to membranous organelles, less is known about how non-membrane bound organelle fusion and fission is maintained. cNap1 intra-organelle assemblies at centriole contacts have a number of the characteristics used to define membrane-membrane contact sites [39], including dedicated tethering machinery that does not induce full fusion of the rest of the organelle. In this regard it is interesting to note that most membrane contact sites are maintained by multiple tethering complexes [40], a feature shared by centrosomes that have both rootletin and cNap1-based tethering.

Previous data has shown that cNap1 anchors rootlets to centrioles, since cNap1 disruption prevents rootlet formation at centrosomes [3,9,26,29]. A previous cyro-electron tomography study also described an “amorphous density”, between centriole pairs and partially inside the centriolar lumen, from which fibres emerge [41]. Consistent with these observations, cNap1 condensates bind specifically to rootletin fibre termini (**Figs. 1** and **3**). One interpretation of this observation is that cNap1 condensates create a phase-separated environment that promotes rootlet fibre anchoring or nucleation at the proximal centriole. This model is reminiscent of others proposed for the pericentriolar material - a different centriole protein coat that has also been suggested to phase separate to nucleate microtubules [42,43].

Overall charge is known to regulate cNap1 oligomerisation, through multisite phosphorylation from the kinase Nek2 [33]. Since multivalent charge-charge interactions are known to regulate condensate formation [44], an interesting future direction of investigation could be to determine whether phosphorylation dependent cNap1 condensate formation and rootlet end-binding control its centrosomal functions.

The data here provide a framework to understand the effects of cNap1 disease-causing mutations in the future. cNap1 R188 is severely truncated, disrupts centrosome cohesion as a dominant negative, and forms condensates with altered material properties (**Fig. 4** and **5**), illustrating how the effects of other mutations may be rationalised.

cNap1 is not conserved throughout the Animalia kingdom, in contrast to its paralog rootletin [5]. The organelle paralogy hypothesis suggests that paralogous duplication is a mechanism for the diversification of membrane-bound organellar function during evolution [45]. This suggests the untested hypothesis that cNap1 has evolved to impart additional centrosomal functionality to mammals.

In conclusion, this work suggests a model of centrosome cohesion using dynamic cNap1 assemblies, and identifies a new intra-organelle contact site. More generally, this provides insight into how a non-membrane bound organelle forms dynamic organelle-organelle contacts within the cellular interior.

## Supporting information

**Fig. S1.**
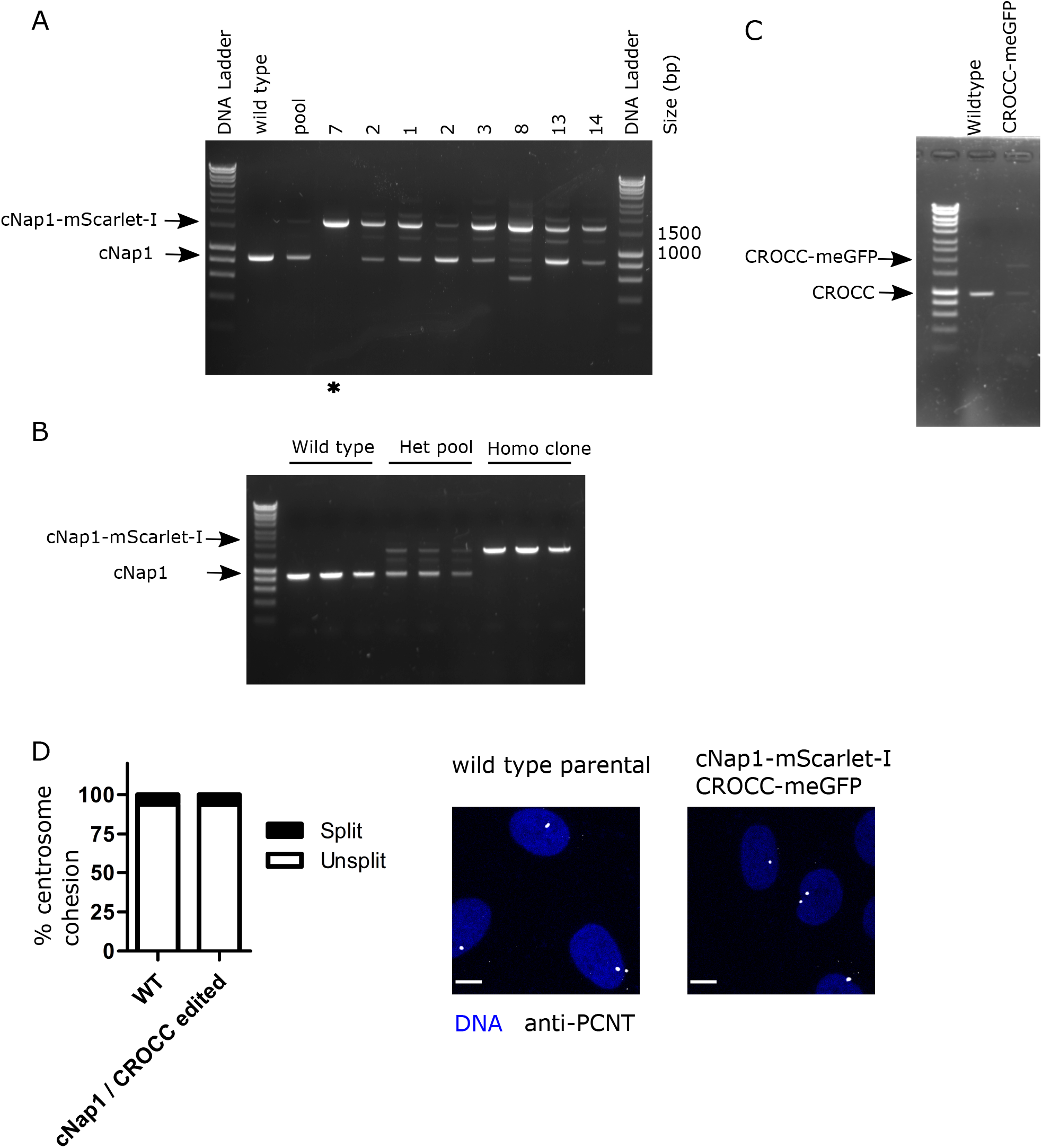
Construction and validation of endogenously tagged cNap1-mScarlet-I and CROCC-meGFP in U2OS cells. (**A**) DNA gel showing junction PCR screening of genomic DNA, for insertion of mScarlet-I at the C-terminus of cNap1. Clone 7 was selected since it is homozygous for cNap1-mScarlet-I. The selected clone is indicated by *. (**B**) Junction PCR of genomic DNA, screening for insertion of mScarlet-I at the C-terminus of cNap1. This shows a comparison of clone 7 with a heterozygous pool. Lanes are loaded in triplicate to exclude the possibility of lane-to-lane variability. (**C**) Junction PCR of genomic DNA, screening for insertion of meGFP at the C-terminus of CROCC. (**D**) Centrosome cohesion in wild type and cNap1-mScarlet-I / rootletin-meGFP cells, assessed by immunofluorescent staining of centrosomes with anti-PCNT antibody in populations of cells. Centrosomes were classed as split if two PCNT positive foci were present and separated by more than 1.6 μm. The data is the mean of three experiments. The images show maximum intensity projections of confocal Airyscan z-stacks. Scale 10 μm.

**Fig. S2.**
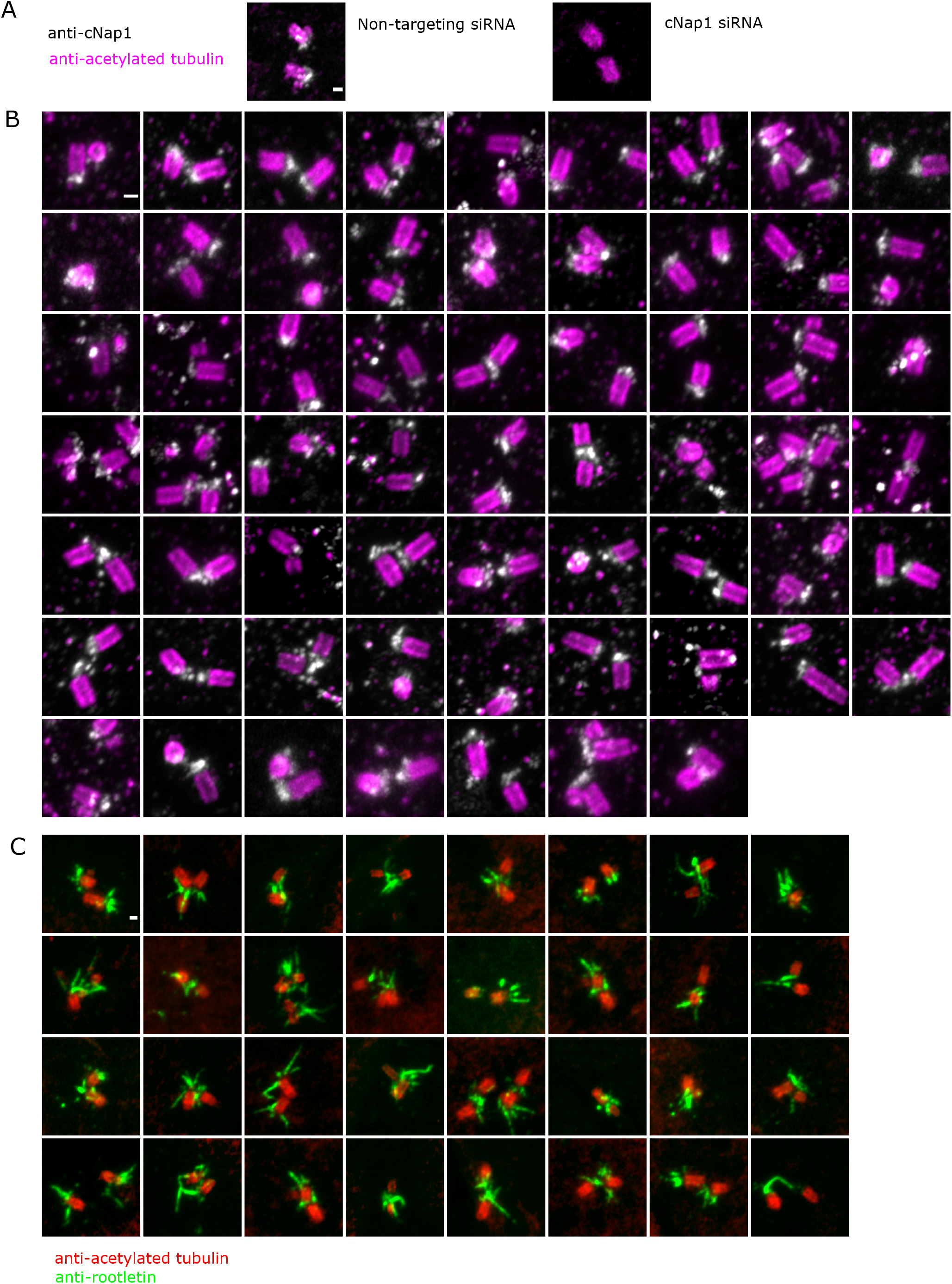
U-ExM of centrioles and cNap1. (**A**) Validation of anti-cNap1 U-ExM staining with siRNA. Cells were treated with either siRNA targeting cNap1 (left panel), or non-targeting siRNA (right panel), and then processed identically for U-ExM. Scale 200 nm. (**B**) U-ExM expanded cells stained with anti-cNap1 (grey) and anti-acetylated tubulin (magenta). Each image is a different cell. Maximum intensity z-projections are shown. Scale 200 nm. (**C**) U-ExM expanded cells stained with anti-rootletin (green) and anti-acetylated tubulin (red). Each image is a different cell. Scale 200 nm.

**Fig. S3.**
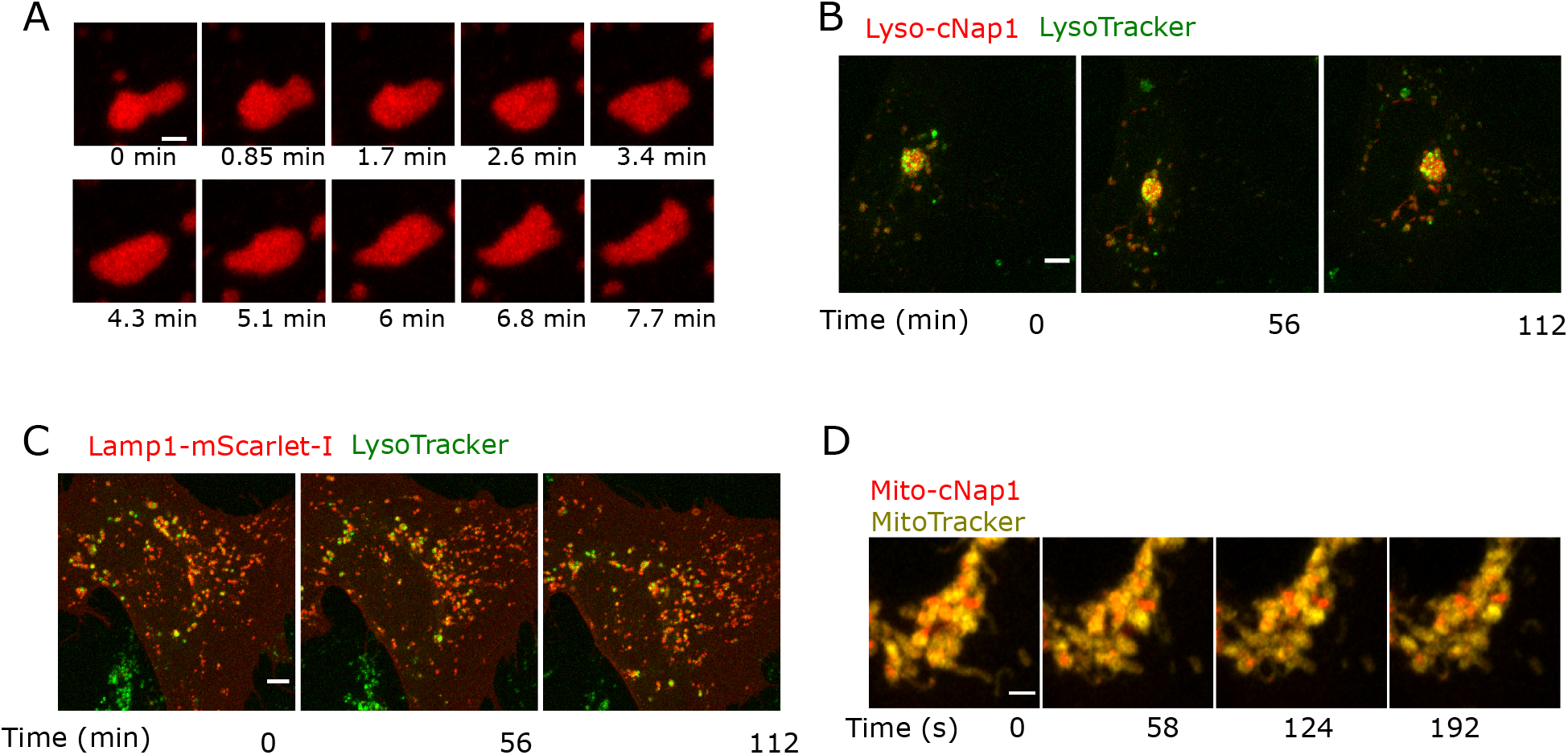
In vivo behaviour of mScarlet-I-cNap1 condensates. (**A**) Live-cell time-lapse imaging of a single cytoplasmic mScarlet-I-cNap1 condensate over mins, showing viscous liquid-like shape changes over time. Scale 1.5 μm. (**B**) Dual colour time-lapse imaging of lyso-cNap1 (red) and LysoTracker (green). Scale 5 μm. (**C**) Dual colour time-lapse imaging of Lamp1-mScarlet-I (red; right panel) and LysoTracker (green). Scale 5 μm. (**D**) Dual colour time-lapse imaging of mito-cNap1 (red) and MitoTracker (yellow). Scale 2 μm.

## Supplementary movies

**S1 Movie**. Time-lapse Airyscan imaging of endogenous cNap1-mScarlet-I in U2OS cells at one min intervals, for a total time of 30 mins, related to **Fig. 1 B**. A sum projection of a z-stack is shown.

**S2 Movie**. Time-lapse Airyscan imaging of endogenous rootletin-meGFP and cNap1-mScarlet-I at 12 s intervals in U2OS cells, related to **Fig. 1 C**. A sum projection of a z-stack is shown.

**S3 Movie**. Time-lapse Airyscan imaging of endogenous rootletin-meGFP and cNap1-mScarlet-I at 12 s intervals in U2OS cells, related to **Fig. 1 D**. A sum projection of a z-stack is shown.

**S4 Movie**. Formation of mScarlet-I-cNap1 condensates in the cytoplasm after transfection. Maximum intensity projections at 30 min time intervals in U2OS cells. Scale 10 μm.

**S5 Movie**. Viscous behaviour of mScarlet-I-cNap1, taken at 30 min intervals in U2OS cells.

**S6 Movie**. Coalescence of mScarlet-I-cNap1 condensates. Time frames are taken at three min intervals, related to **Fig. 3 C**. Scale 2 μm.

**S7 Movie**. Time-lapse imaging of eGFP-rootletin and mScarlet-I-cNap1 at 0.5 h intervals. Maximum intensity projections are shown. Scale 10 μm.

**S8 Movie**. Time-lapse imaging of eGFP-rootletin and mScarlet-I-cNap1 at 5 min intervals, related to **Fig. 3 F**.

**S9 Movie**. Time-lapse Airyscan imaging of LysoTracker (green) and lyso-cNap1 (red) at 14 min intervals, related to **Fig. 5 F**. A maximum intensity z-projection is shown. Scale 5 μm.

**S10 Movie**. Time-lapse Airyscan imaging of LysoTracker (green) and Lamp1-mScarlet-I (red) at 14 min intervals. A maximum intensity z-projection is shown. Scale 5 μm.

**S11 Movie**. Time-lapse Airyscan imaging of mito-cNap1 (red) and MitoTracker (yellow) at 7 second intervals. A sum z-projection is shown. Scale 2 μm.

**S12 Movie**. Time-lapse Airyscan imaging of R188 mScarlet-I-cNap1 condensates, showing reduced fusogenic capability relative to wild type. Images are taken at 3 min intervals. Image show maximum intensity projections. Scale 4 μm.

## Materials and Methods

### Cell culture and chemicals

U2OS cells were obtained directly from the American Type Culture Collection (ATCC HTB-96). U2OS and HeLa Kyoto cell lines were grown in Dulbecco’s modified Eagle’s Medium (DMEM) supplemented with 10% fetal calf serum, Glutamax, and 100 μg/ml penicillin/streptomycin and maintained at 37 °C with 5% CO_2_. Cell lines were confirmed as mycoplasma free. All tissue culture reagents were purchased from Sigma-Aldrich unless otherwise stated. DNA transfection was with lipofectamine 3000 (Invitrogen) or jetPRIME (Polyplus) according to the manufacturer’s instructions.

### CRISPR cas9-mediated genome editing

Endogenously tagged cNap1-mScarlet-I and rootletin-meGFP (CROCC-meGFP) U2OS cells were produced with the methods described in [46]. Donor plasmids consisted of two 800 bp homology arms surrounding the C-terminus of either the *cNap1* or *CROCC* genomic reference sequence. These arms were inserted into plasmids such that they flank mScarlet-I or meGFP coding sequence. The cNap1 donor plasmid was purchased from ThermoFisher GeneArt. The *CROCC* donor plasmid was constructed in-house by In-fusion cloning (see molecular cloning for in-fusion methods).

sgRNA sequences were designed in Benchling software, selecting optimal on and off target activity as close to the target site as possible. Guide RNAs did not target the donor plasmid. Guide RNA sequences (5’—3’) for *cNap1* were: TCCAGGTAGCAGCCACAGCC (Strand 1), CTGTGGCTGCTACCTGGAGG (Strand -1), TCCTGGCTGTGGCTGCTACC (Strand -1). Guide RNA sequences (5’—3’) for *CROCC* against the +ve strand were as follows (5’—3’): CCAGCAGGAGCTCATTTCTC, CCAGAGAAATGAGCTCCTGC, and CAGGAGCTCATTTCTCTGGG.

Guide RNA sequences were cloned into pSpCas9(BB)-2A-Puro (PX459) (Addgene plasmid 48139) for expression. Guide RNA and the donor plasmid were co-transfected. After a week, single cells were sorted for mScarlet-I or meGFP positivity and grown as clones for PCR screening. Genomic insertion of fluorescent proteins was screened by overlapping genomic PCR. PCR primers were designed either side of the insertion site in clone manager suite, ensuring no false priming. PCR primer annealing temperature was optimised across a temperature gradient before screening clones. For CROCC-meGFP, primer sequences were as follows 5’-3’: GGCTTGGATCTAAGGAGG and GGCTGGCCTTACCTTCCCTT. For cNap1-mScarlet-I primer sequences were as follows 5’-3’: GATTCGTGTATGTGGTAGAG and CTATCACAGTGCATGGTGTA. Tag insertion was detected on the basis of PCR product size, ~700 bps larger with fluorescent protein insertion. PCR for screening was with DREAMtaq (Invitrogen) according to the manufacturer’s instructions, run with Hyperladder 1 DNA marker (Bioline). Selected clones were confirmed to have centrosomal localisation as expected by Airyscan confocal imaging, concurring with previously reported antibody staining. Cell lines were also validated by removing fluorescent signal using siRNA-mediated knockdown of either cNap1 or rootletin, using the methods described in see siRNA methods section.

### Airyscan and confocal imaging

All images except those in **Fig. 3A** were acquired on a Carl Zeiss LSM 880 Airyscan confocal laser scanning microscope, controlled by Zen Black software. Lenses used were 100x NA 1.46 oil, 63x NA 1.4 oil, 40x NA 1.3 oil and 40x NA 1.2 water immersion objectives, with optimised cover glass correction where possible. Airyscan images were acquired in SR mode and processing was performed with automatic settings in Zen Black. Pixel size and Z-slice size were optimised, depending on the scan area of each experiment (which was variable), using the optimal function in Zen. Other imaging parameters including scan speed and image averaging were variable for each experiment, but never changed between comparative samples. Laser power was adjusted to minimise bleaching and cellular toxicity in live-cell experiments. Detector gain was adjusted to ensure pixel intensities were never saturated or clipped. The median (± median absolute deviation) lateral and axial resolution of the system was measured at 198 ± 7.5 nm and 913 ± 50 nm (full width at half-maximum), respectively, through imaging of a sub resolution fluorescent bead. Brightness and contrast were adjusted on images for display purposes, but never unequally between comparative samples.

### Live-cell time-lapse imaging and FRAP

Cells were imaged in L15 CO_2_-independent medium at 37 °C. Cell health in the conditions used was optimal since cell growth continued. For live-cell timelapse, image size and frequency of acquisition varied depending on the timescale of events observed and scan area. For high resolution imaging of centrosomes a 100x NA 1.46 Oil lens was used. Autofocus was at every timepoint using Zeiss definite focus autofocus system. A stage top Z-piezo was used for high-speed z-stack imaging. FRAP was performed essentially as described previously [23]. Selected image regions were bleached with a 561-laser line at 100% for the minimum time required to cause approximately 50% fluorescence loss. The bleach duration was constant in all samples. Cells were imaged in a single z-plane following bleaching, at ~0.7 s intervals. Analysis was in Microsoft Excel and GraphPad Prism. Images were background subtracted. With the imaging conditions used, bleaching was minimal, as determined by running the experiment with identical settings except for the FRAP bleach, and measuring the change in intensity.

### Expansion microscopy

U-ExM was as described in [31,32,47]. Cells were seeded on 12 mm coverslips overnight, before fixation for 5 hrs in humid conditions at 37 °C in 1.4% / 2% formaldehyde (F8775 Sigma) / acrylamide (A4058 Sigma). Gelation was in U-ExM monomer solution, consisting of 23% w/v sodium acrylate (408220 Sigma), 10% w/v acrylamide and 0.1% w/v N,N’-methylenbisacylamide in PBS. 0.5% tetramethylethylenediamine (17919, ThermoFisher) and 0.5% ammonium persulfate (17874, ThermoFisher) were added to the monomer solution directly prior to gelation, with the samples on ice. Gelation was for five mins on ice and one h at 37 °C, in a humid chamber. Denaturation was in U-ExM denaturation buffer (200 mM SDS, 200 mM NaCl, 50 mM Tris-BASE in ddH_2_O) at 95 °C for 90 mins [47]. Gels were expanded overnight at room temperature in ddH_2_O prior to and post antibody incubation. Primary antibody labelling was either overnight at 4 °C or 3 hrs at 37 °C in 2 % BSA-PBS at 1:250 dilution. Secondary antibody labelling was at 37 °C in 2 % BSA-PBS at 1:500 dilution for 2.5 hrs. Gels were washed with PBS 0.1 % triton-X after both antibody staining steps. Nuclei were labelled with Hoescht 33342 dye in the final wash step. Gels were mounted in Ibidi μ-slide 2 well glass bottom #1.5 dishes (80286), which were pre-treated in poly-L-lysine or poly-D-lysine and imaged using Airyscan imaging. Primary antibodies used were mouse anti-acetylated tubulin (Sigma Aldrich, T7451), rabbit anti-rootletin (Novus Biologicals, NBP1-80820), rabbit anti-cNap1 (Proteintech, 14498-1-AP), mouse anti-γtubulin (GTU-88, Abcam) and mouse anti-centrin 2 (Millipore, 04-1624 20H5). For anti-γtubulin only, cells were fixed in 100% ice-cold methanol for five mins prior to formaldehyde fixation. Secondary antibodies were Alexa 488, Alexa 568 or Atto 565 conjugates. The expansion factor was calculated from measurements of the gel diameter pre and post expansion, and from centriole size in final images when stained with anti-acetylated tubulin, giving values around 4.2 - 4.6. Calculation of the percentage of cells with cNap1 bridges between centrioles was done from three independent experiments, measuring a total of ~80 cells.

### Antibody validation

cNap1 antibody was validated specifically in U-ExM by confirming that signal was removed by siRNA targeting cNap1 in comparison to non-targeting control siRNA (**Fig. S2 A**). Moreover, in standard immunofluorescence, anti-cNap1 staining closely matched cNap1-mScarlet-I fluorescent protein.

### Automated image acquisition and analysis

Images in **Fig. 3 A** were acquired on a Molecular Devices ImageXpress Micro Confocal. Objective cover glass corrections were optimised to scan with Ibidi μ-slide 8-well dishes and a 40x air objective. Multiple z-sections were obtained and then projected using the Molecular Devices “best” function. Images were analysed in Ilastik and CellProfiler software, using a custom-made pipelines. Briefly, cells and cNap1 foci were automatically segmented using pixel-based image classification. Segmented images were further analysed in CellProfiler, using the relate function and to associate cNap1 and cells and therefore count number per cell. Segment shape parameters were calculated, dividing the major and minor axis lengths to obtain the aspect ratio.

### Molecular cloning

DNA constructs were made by In-fusion HD cloning (Clontech) into the vector pcDNA 3.1, according to the manufacturer’s instructions. Briefly, primers containing complementary 15 bp extensions were designed in the TaKaRa Bio In-fusion online design tool. Both the vector and inserts were amplified by PCR with CloneAmp DNA polymerase. Amplified DNA length was verified by agarose gel electrophoresis. In-Fusion ligation was performed on gel extracted DNA using In-Fusion HD enzyme premix in a total volume of 5 μl at 50 °C for 15 mins. Clones were screened by Sanger DNA sequencing, restriction digest and microscopy after transfection into mammalian cells. cNap1 cDNA was originally provided by Andrew Fry (University of Leicester, UK).

### Design of Mito-cNap1, Lyso-cNap1, Golgi-cNap1 and cNap1 truncation constructs

Mito-cNap1 consists of an N-terminal fusion of a pair of mitochondria targeting sequences from cytochrome c oxidase subunit VIII (COX8) [48], separated by a short linker, to give the following amino acids: MSVLTPLLLRGLTGSARRLPVPRAKIHSLPPEGKLGMSVLTPLLLRGLTGSARRLPVPRA. This was fused in frame to either mScarlet-I alone or mScarlet-I-cNap1, therefore forming COX8-mScarlet-I or COX8-mScarlet-I-cNap1. Lyso-cNap1 consists of cNap1-mScarlet-I fused in frame to human lysosomal-associated membrane protein 1 (*LAMP-1*) [49], to create LAMP1-cNap1-mScarlet-I. A negative control consisted of LAMP1-mScarlet-I. Golgi-cNap1 consists of a C-terminal fusion of a GRIP domain [50], consisting of the C-terminal 98 amino acids of Golgin-245, to cNap1-mScarlet-I. This therefore creates cNap1-mScarlet-I-GRIP. A negative control consisted of mScarlet-I-GRIP. cNap1 truncations were made by HD In-fusion cloning from the full-length gene.

### siRNA transfection

siRNA targeting rootletin (gene name *CROCC*) was as previously described by [3], and is as follows: 5′-AAGCCAGTCTAGACAAGGA-3′. This siRNA notably has a strong centrosome splitting phenotype relative to other rootletin-targeting siRNA [3], and was synthesised by Horizon Discovery. Non-targeting negative control siRNA and siRNA targeting cNap1 were ON-TARGET *plus* pools from Horizon Discovery (D-001810-OX and L-012364 respectively). siRNA transfection was with RNAiMAX (Thermo Fisher Scientific), following the manufacturer’s instructions. Briefly, for 96-well transfections, cells were transfection with 25 nM of siRNA, and 0.25 μl lipofectamine per well. Cells were analysed either 48 or 72 hrs after transfection. The efficacy of both cNap1 and rootletin siRNA knockdown was confirmed by loss of fluorescence in cNap1-mScarlet-I and rootletin-meGFP cells.

### Standard immunofluorescence and dye staining

Cells were fixed in either 4% paraformaldehyde in PBS pH 7.4 for 15 min or ice-cold 100% methanol for 5 min. Fixatives were freshly prepared. Paraformaldehyde was quenched in 0.1 M NH_4_Cl in PBS pH 7.4. Cells were permeabilised in 0.1% Triton in PBS and blocked in 3% bovine serum albumen (ThermoFisher Scientific) in PBS. Antibodies used were: mouse anti-gamma tubulin GTU-88 (1:1000 Sigma-Aldrich T6557), rabbit anti-GM130 (1:1000 Abcam, ab52649), rabbit anti-PCNT (1:1000 Abcam ab4448). MitoTracker deep red (ThermoFisher Scientific) was incubated at culture conditions for five mins at 100 nM before replacing with fresh medium for imaging. Lysotracker was used at 75 nM, added directly prior to imaging and kept in the imaging medium.

### Statistical Analysis

Statistical analyses and graphical representations were in GraphPad Prism 5 software. Statistical tests are listed in the figure legends.

## Acknowledgements

This work was funded by a Wellcome Trust Henry Wellcome Fellowship to R.M. (https://wellcome.ac.uk/grant number 100090/12/Z), by the MRC Cancer Unit and the Isaac Newton Trust (grant number 21.23(j)). I thank the Cambridge Institute for Medical Research flow cytometry core facility for cell sorting. Suzan Ber Esposito, Annie Howitt, Alessandro Esposito and Andrew Fry kindly provided cDNA template plasmids. I thank Helen Pickersgill, Marco Sciacovelli, Dylan Ryan and Amy Emery for constructive comments on the draft manuscript.

## Author contributions

All aspects: R.M.

